# Modelling Niemann-Pick disease type C in a human haploid cell line allows for patient variant characterization and clinical interpretation

**DOI:** 10.1101/801019

**Authors:** Steven Erwood, Reid A. Brewer, Teija M.I. Bily, Eleonora Maino, Liangchi Zhou, Ronald D. Cohn, Evgueni A. Ivakine

## Abstract

The accurate clinical interpretation of human sequence variation is foundational to personalized medicine. This remains a pressing challenge, however, as genome sequencing becomes routine and new functionally undefined variants rapidly accumulate. Here, we describe a platform for the rapid generation, characterization and interpretation of genomic variants in haploid cells focusing on Niemann-Pick disease type C (NPC) as an example. NPC is a fatal neurodegenerative disorder characterized by a lysosomal accumulation of unesterified cholesterol and glycolipids. In 95% of cases, NPC is caused by mutations in the *NPC1* gene, where over 200 unique disease-causing variants have been reported to date. Furthermore, the majority of patients with NPC are compound heterozygotes that often carry at least one private mutation, presenting a challenge for the characterization and classification of individual variants. Here, we have developed the first haploid cell model of NPC. This haploid cell model recapitulates the primary biochemical and molecular phenotypes typically found in patient-derived fibroblasts, illustrating its utility in modelling NPC. Additionally, we demonstrate the power of CRISPR-Cas9-mediated base editing in quickly and efficiently generating haploid cell models of individual patient variants in NPC. These models provide a platform for understanding the disease mechanisms underlying individual *NPC1* variants while allowing for definitive clinical variant interpretation for NPC. While this study has focused on modelling NPC, the outlined approach could be translated widely and applied to a variety of genetic disorders or understanding the pathogenicity of somatic mutations in cancer.

## Introduction

Niemann-Pick disease type C is a rare autosomal recessive lysosomal storage disorder affecting 1 in 90,000 individuals (Vanier 2010; Wassif et al. 2016). In 95% of cases, NPC is caused by mutations in the gene *NPC1*, which is required for the proper transport of sterols from the lysosome to other subcellular compartments (Vanier 2010). While NPC is a clinically heterogeneous disorder with symptoms ranging from hepatosplenomegaly to ataxia and seizures, the disease is defined by fatally progressive neurodegeneration (Patterson et al. 2013; Vanier 2010). These symptoms are caused by the intracellular accumulation of cholesterol and glycolipids within late endosomes and lysosomes (Wojtanik and Liscum 2003; Ory 2000). This accumulation is easily visualized in patient-derived fibroblasts using a fluorescent dye called filipin which is used as a primary assay in the diagnosis of Niemann-Pick disease type C (McKay Bounford and Gissen 2014).

More than 200 disease-causing mutations have been identified in *NPC1* which define a heterogeneous mutational spectrum that includes missense and nonsense mutations, small duplication, deletion and insertion mutations, and splice-site mutations (Scott and Ioannou 2004; Park et al. 2003; Fernandez-Valero et al. 2005; Millat et al. 2001; Tarugi et al. 2002). The primary source material used to understand NPC pathology in humans is patient-derived fibroblasts (Gelsthorpe et al. 2008; Greer et al. 1999; Millat et al. 2001; Rauniyar et al. 2015; Yamamoto et al. 2004; Zampieri et al. 2012). The majority of patients with NPC, however, present as compound heterozygotes that often harbour at least one private mutation. This presents a challenge in understanding the molecular mechanisms of disease underlying individual *NPC1* variants, leaving most documented mutations as variants of uncertain significance. Further complicating variant interpretation, it has been demonstrated that variant pathogenicity is contingent on level of expression. Specifically, certain variants that are pathogenic at physiologically relevant expression levels can rescue disease phenotypes when artificially overexpressed (Gelsthorpe et al. 2008; Zampieri et al. 2012).

The advent of CRISPR-Cas9-based genome editing has allowed for modifications to genomes with a precision and efficiency unparalleled by previous technologies (Mali, Esvelt, and Church 2013). In brief, CRISPR-Cas9-based genome editing relies on a guide RNA programmable bacterial endonuclease, Cas9, to induce a targeted DNA double-stranded break (DSB). In the absence of a repair template, this break is predominantly repaired by non-homologous end joining (NHEJ) which is stochastic and leads to small insertions or deletions (Cho et al. 2013; Jinek et al. 2012; Mali et al. 2013). Typically, even when a repair template is exogenously supplied, NHEJ is responsible for the majority of genome editing outcomes with CRISPR-Cas9, making the establishment of models with specifically designed modifications inefficient. Recently, this challenge has been addressed with the introduction of CRISPR-Cas9-mediated base editing, which utilizes a nucleobase deaminase enzyme fused to a catalytically impaired Cas9 enzyme capable of inducing only single-stranded breaks (Rees and Liu 2018). These nucleobase deaminase enzymes, APOBEC1 and TadA for cytosine and adenine base editing, respectively, operate on single-stranded DNA (ssDNA), exclusively (Gaudelli et al. 2017; Komor et al. 2016). Similar to traditional CRISPR-Cas9-based genome editing, this fusion protein can be targeted to a guide RNA specified genomic locus. When the guide RNA binds to its target sequence, the complementary strand is displaced, becoming available for modification by the deaminase enzyme (Nishimasu et al. 2014). In practice, only a portion of the displaced “R loop” is prone to deamination with the current generation of CRISPR-Cas9 base editors, corresponding to an approximately 5 base pair editing window located 13-17 base pairs upstream from the protospacer-adjacent motif sequence (PAM) (Gaudelli et al. 2017; Komor et al. 2016; Rees et al. 2017). Deamination of cytosine produces uridine which base pairs as thymidine, while deamination of adenosine produces inosine which has base pairing preferences equivalent to guanosine (Yasui et al. 2008). The single-stranded nick produced on the unedited strand by the Cas9 enzyme then induces endogenous DNA repair pathways that will utilize the edited strand as a template, effectuating either a C·G to T·A or an A·T to G·C base pair transition.

Here, we demonstrate that by CRISPR-Cas9-mediated *NPC1* gene editing, the HAP1 cell line, a human near-haploid cell line, can serve as an effective model of Niemann-Pick disease type C. By CRISPR-Cas9-based gene disruption, we show that *NPC1*-deficient HAP1 cells exhibit the biochemical hallmarks of the disease. Furthermore, we demonstrate that CRISPR-Cas9-mediated base editing allows for the rapid and efficient generation of haploid models of unique *NPC1* variants. These models allow for the interrogation of individual variants in isolation, which was previously infeasible for the vast majority of documented variants. Additionally, we demonstrate the utility of our haploid models of *NPC1* variants in clinical variant interpretation.

## Results

### Generation and characterization of an NPC1-deficient near-haploid cell line

As NPC is an autosomal recessive disorder, cellular disease modelling requires the disruption of each allele present in the target cell type. This presents a challenge given the diploid or often aneuploid nature of typical human cell lines, especially if uniform allele disruption is desired. HAP1 cells, however, are a near-haploid human cell line containing a single copy of each chromosome, with the exception of a heterozygous fragment of chromosome 15, making them an excellent system for loss-of-function disease modelling (Carette et al. 2011). We used CRISPR-Cas9-mediated gene targeting to generate NPC1-deficient HAP1 cells. To disrupt *NPC1* expression, we selected several single guide RNAs (sgRNAs) with minimal computationally predicted off-target activity which target exon 21 of the *NPC1* locus. These sgRNAs were cloned into plasmids allowing co-expression with *Streptococcus pyogenes* Cas9 (SpCas9) and tested for editing efficiency. The two most highly active sgRNAs were transfected separately into wildtype HAP1 cells (Supplemental Table S1, Fig. 1A). Following 72 hours of antibiotic selection to enrich for successfully transfected cells, isogenic clones were isolated by limited dilution and screened by Sanger sequencing for locus disruption. Out of fifteen clones screened, six isogenic cell clones were identified with unique mutations in the targeted locus and three clones were carried forward for further characterization. A twenty-eight base pair deletion was detected in the first clone, [NCBI Reference Sequence: NG_012795.1 *NPC1*: g.54927_54954del], resulting in a frame-shift and the formation of a premature stop codon six amino acids downstream from the deletion site (Fig. 1B). The second clone, [NG_012795.1 *NPC1*: g.54902insA], harboured an insertion of an adenine nucleotide at the predicted double-stranded break site, resulting in a frameshift and the formation of a premature stop codon four amino acids downstream from the insertion (Fig. 1C). The third clone, [NG_012795.1 *NPC1*: g.54899_54904del], contained an in-frame six base pair deletion (Fig. 1D). To assess whether these mutations were sufficient to disrupt NPC1 expression, we performed a western blot for NPC1 protein. Both clones with frameshift mutations exhibited a complete absence of NPC1 protein, while the third clone showed residual protein expression and appeared to run as a doublet, with a second band at a slightly lower molecular weight (Fig. 1E). For the first two clones, this indicates that both frameshift mutations are sufficient in generating null *NPC1* alleles. In the third clone, the doublet staining of NPC1 likely indicates a heterogeneously glycosylated protein product, a phenomenon which has been previously reported for a variety of NPC variants (Watari et al. 1999; Nakasone et al. 2014; Zampieri et al. 2012), and the reduced expression indicates that perturbations to the *NPC1* locus, in spite of a preserved reading frame, can negatively influence protein expression. Our three cell clones, with their disrupted NPC1 protein expression, represent the first haploid cell models of Niemann-Pick disease type C.

**Figure 1.**
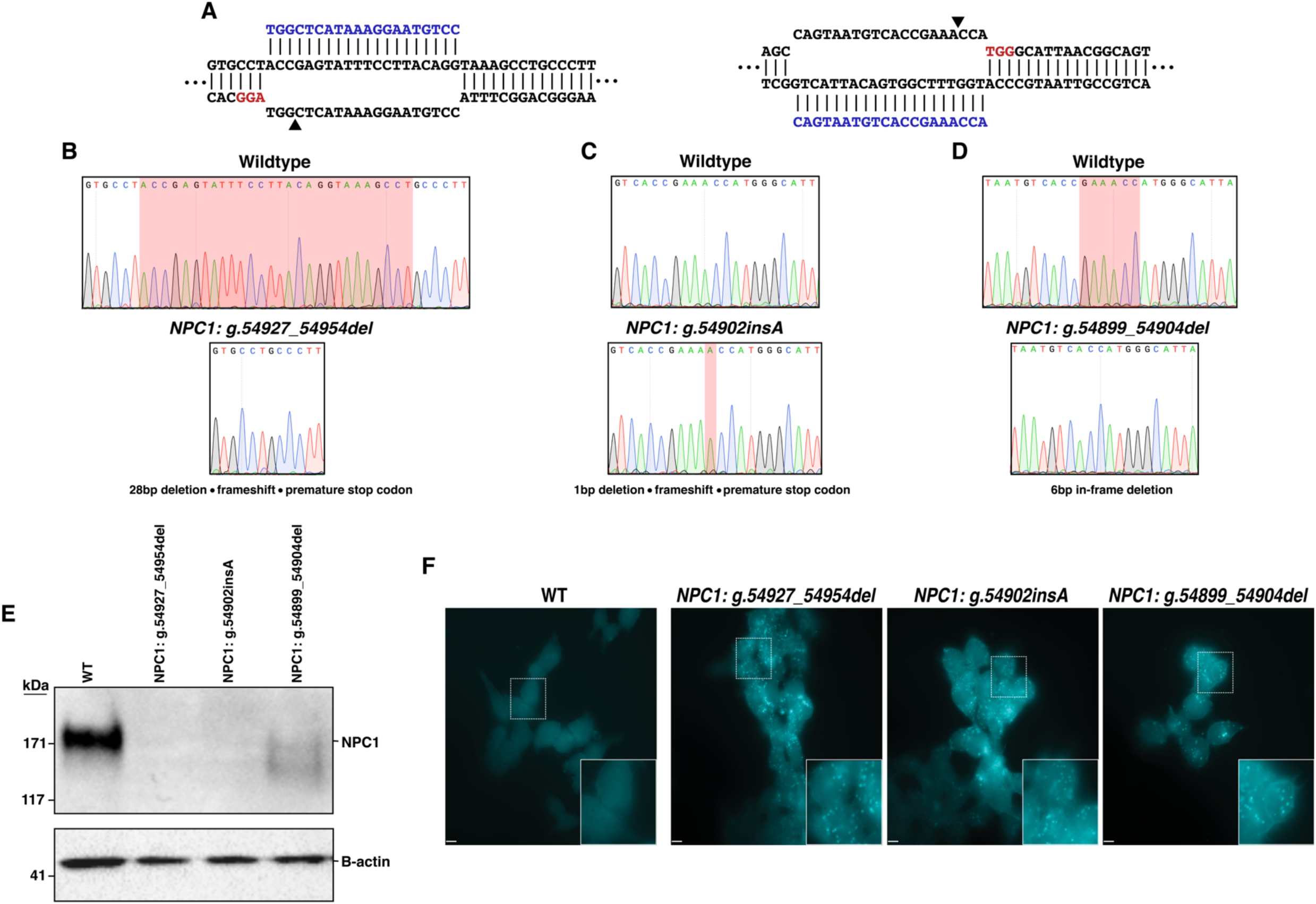
Generation and characterization of three haploid models of Niemann-Pick disease type C. (A) Diagrams illustrating the two targeted sites in *NPC1*. Arrowheads indicate predicted DSB site. (B, C,D) sequencing chromatographs showing wildtype *NPC1*, top, and the specific disruption in each isogenic edited cell clone, bottom. (B, D) Red highlighted region indicates the locations of the deletions in clones. (C) Red highlighted region indicates the position of the insertion in the edited clone. (E) Western blot analysis from total protein lysate from wildtype HAP1 cells and the three edited cell clones ting absent or reduced NPC1 protein expression. B-actin expression was used as a loading control. (F) Filipin staining reveals deposits of intracellular cholesterol in edited cells that are absent in wildtype White dashed-bordered box has been enlarged twofold and inset at bottom right. Scale bars:

A hallmark biochemical feature of NPC pathology is the accumulation of unesterified cholesterol and glycolipids within late endosomes and lysosomes. Presently, the demonstration of defective intracellular cholesterol transport and homeostasis is considered the most definitive functional diagnostic test for NPC (McKay Bounford and Gissen 2014). This defect is readily visualized in NPC patient fibroblasts by staining with the fluorescent compound filipin, which stains unesterified cholesterol deposits (Supplemental Fig. S1). We assessed whether the NPC1-deficient HAP1 cells displayed a similar biochemical phenotype using filipin staining. All three cell clones display distinct foci of intracellular filipin staining that are absent in unedited HAP1 cells, indicative of impaired trafficking of unesterified cholesterol in the NPC1-deficient HAP1 cells (Fig. 1F). The development of disease-relevant pathology in NPC1-deficient HAP1 cell clones demonstrates the potential of these cells in understanding disease mechanisms in NPC.

### Modelling NPC1 variants in a haploid cell model using CRISPR-Cas9-mediated base editing

After demonstrating the effectiveness of HAP1 cells in recapitulating a primary cellular phenotype of NPC, we sought to investigate the feasibility of modelling individual *NPC1* variants using HAP1 cells. To do this, we elected to use CRISPR-Cas9-mediated base editing. CRISPR-Cas9-mediated base editing technologies are capable of targeted single-nucleotide transitions within a designated editing-window upstream from the PAM sequence (Gaudelli et al. 2017; Komor et al. 2016). We selected mutations that span the NPC1 protein and that are representative of the *NPC1* mutation spectrum, including missense, nonsense and synonymous mutations and a splice site mutation. We selected variants that have or have not been previously documented in clinical databases, and variants of both known and unknown pathogenicity. The nineteen variants modelled are documented in Table 1. While we focused on the C-terminal luminal loop domain, spanning residues 855-1098 of NPC1, where 45% of NPC patient mutations occur (Greer et al. 1999; Li et al. 2017), we selected sgRNAs to establish at least one mutation in each of five functional protein domains (Davies and Ioannou 2000). These sgRNA cassettes were cloned into a U6-driven expression vector and individually co-transfected into wildtype HAP1 cells alongside either a Cas9 cytosine or adenine base editor plasmid (Zafra et al. 2018; Koblan et al. 2018; Huang et al. 2019; Nishimasu et al. 2018). Following antibiotic selection, transfected cells were subject to limited dilution to isolate isogenic clones. In each case, editing was apparent in a bulk population, ranging from 10%-48%, (Supplemental Fig. S2). When clones were individually screened by Sanger sequencing by analyzing the genomic sequence ranging from at least 100bp both up- and downstream of the sgRNA binding site, between 8%-60% of isogenic clones were positively edited (Table 1). Our system for model generation resulted in clonal isolation in just over two weeks (Fig. 2A). The editing in all but two of the nineteen variants isolated was contained to a single codon. During the generation of the NPC1 p.R1077X variant, the editing window contained a second cytosine adjacent to the targeted cytosine, and in our screened clones, we were only able to identify clones where both bases were edited. The secondary mutation, however, is a silent mutation where the adjacent histidine codon has been changed from TAC to TAT, resulting in an NPC1 p.Y1076=/R1077X cell line (hereafter referred to as NPC1 p.R1077X) (Supplemental Fig. S3). Similarly, when isolating the NPC1 p.I1061T variant, an adjacent adenine two bases upstream was uniformly targeted in all edited clones, resulting in a secondary silent mutation and an NPC1 p.L1060=/I1061T cell line (hereafter referred to as NPC1 p.I1061T). For the rest of the *NPC1* variants, however, multiple clones were isolated with editing contained to a single codon (Table 1, Supplemental Fig. S3). These data demonstrate that both CRISPR-Cas9-mediated cytosine and adenine base editing are highly efficient in HAP1 cell model generation, providing a viable solution to the documented poor efficiency of introducing single-nucleotide variants (SNVs) by typical CRISPR-Cas9-based homology-directed repair, which is particularly inefficient in HAP1 cells (Findlay et al. 2018).

**Table 1.** Summary of NPC1 variants modelled.

| Variant | Editor | Nucleotide change | ClinVar annotation | Functional outcome observed | Positive Clones (%) |
| --- | --- | --- | --- | --- | --- |
| NPC1 p.R4H | SpCas9-NG Target-AID | CGC > CAC | *Undocumented* | Benign | 8 |
| NPC1 p.F402P | SpCas9 ABEmax | TTC > CCC | *Undocumented* | Pathogenic | 15 |
| NPC1 p.F402S | SpCas9 ABEmax | TTC > TCC | *Undocumented* | Pathogenic | 43 |
| NPC1 p.E406G | SpCas9 ABEmax | GAG > GGG | *Undocumented* | Pathogenic | 33 |
| NPC1 p.Y634Y | SpCas9 ABEmax | TAT > TAC | *Undocumented* | Benign | 10 |
| NPC1 p.Y634C | SpCas9 ABEmax | TAT > TGT | *Variant of uncertain significance* | Pathogenic | 10 |
| NPC1 p.I635T | SpCas9 ABEmax | ATT > ACT | *Undocumented* | Benign | 15 |
| NPC1 p.G765K | SpCas9-NG Target-AID | GGA > AAA | *Undocumented* | Pathogenic | 14 |
| NPC1 p.L785S | NG-Cas9 ABEmax | TTA > TCA | *Undocumented* | Pathogenic | 60 |
| NPC1 p.D898N | SpCas9 CBE | GAC > AAC | *Variant of uncertain significance* | Pathogenic | 50 |
| NPC1 p.D945N | SpCas9 CBE | GAT > AAT | *Variant of uncertain significance* | Pathogenic | 43 |
| NPC1 p.L1060P | NG-Cas9 ABEmax | CTT > CCC | *Undocumented* | Pathogenic | 60 |
| NPC1 p.L1060=, I1061T | VRQR-SpCas9 ABEmax | CTT ATA > CTC ACA | *Pathogenic* | Pathogenic | 8 |
| NPC1 p.Y1076=, R1077X | SpCas9 CBE | TAC CGA > TAT TGA | *Pathogenic* | Pathogenic | 21 |
| NPC1 p.D1097N | SpCas9 CBE | GAC > AAC | *Likely pathogenic* | Pathogenic | 20 |
| NPC1 p.V1158= | SpCas9 CBE | GTG > GTA | *Undocumented* | Benign | 21 |
| NPC1 p.M1159T | SpCas9 ABEmax | ATG > ACG | *Undocumented* | Benign | 27 |
| NPC1 c.3591+2 T>C | SpCas9 ABEmax | GTG > GCG | *Likely pathogenic* | Pathogenic | 25 |
| NPC1 p.Y1267C | VRQR-Cas9 ABEmax | TAC > TGC | *Undocumented* | Benign | 33 |

Table 1. Summary of NPC1 variants modelled.
| Variant | Editor | Nucleotide change | ClinVar annotation | Functional outcome observed | Positive Clones (%) |
| --- | --- | --- | --- | --- | --- |
| NPC1 p.R4H | SpCas9-NG Target-AID | CGC > CAC | Undocumented | Benign | 8 |
| NPC1 p.F402P | SpCas9 ABEmax | TTC > CCC | Undocumented | Pathogenic | 15 |
| NPC1 p.F402S | SpCas9 ABEmax | TTC > TCC | Undocumented | Pathogenic | 43 |
| NPC1 p.E406G | SpCas9 ABEmax | GAG > GGG | Undocumented | Pathogenic | 33 |
| NPC1 p.Y634Y | SpCas9 ABEmax | TAT > TAC | Undocumented | Benign | 10 |
| NPC1 p.Y634C | SpCas9 ABEmax | TAT > TGT | Variant of uncertain significance | Pathogenic | 10 |
| NPC1 p.I635T | SpCas9 ABEmax | ATT > ACT | Undocumented | Benign | 15 |
| NPC1 p.G765K | SpCas9-NG Target-AID | GGA > AAA | Undocumented | Pathogenic | 14 |
| NPC1 p.L785S | NG-Cas9 ABEmax | TTA > TCA | Undocumented | Pathogenic | 60 |
| NPC1 p.D898N | SpCas9 CBE | GAC > AAC | Variant of uncertain significance | Pathogenic | 50 |
| NPC1 p.D945N | SpCas9 CBE | GAT > AAT | Variant of uncertain significance | Pathogenic | 43 |
| NPC1 p.L1060P | NG-Cas9 ABEmax | CTT > CCC | Undocumented | Pathogenic | 60 |
| NPC1 p.L1060=, I1061T | VRQR-SpCas9 ABEmax | CTT ATA > CTC ACA | Pathogenic | Pathogenic | 8 |
| NPC1 p.Y1076=, R1077X | SpCas9 CBE | TAC CGA > TAT TGA | Pathogenic | Pathogenic | 21 |
| NPC1 p.D1097N | SpCas9 CBE | GAC > AAC | Likely pathogenic | Pathogenic | 20 |
| NPC1 p.V1158= | SpCas9 CBE | GTG > GTA | Undocumented | Benign | 21 |
| NPC1 p.M1159T | SpCas9 ABEmax | ATG > ACG | Undocumented | Benign | 27 |
| NPC1 c.3591+2 T>C | SpCas9 ABEmax | GTG > GCG | Likely pathogenic | Pathogenic | 25 |
| NPC1 p.Y1267C | VRQR-Cas9 ABEmax | TAC > TGC | Undocumented | Benign | 33 |

**Figure 2.**
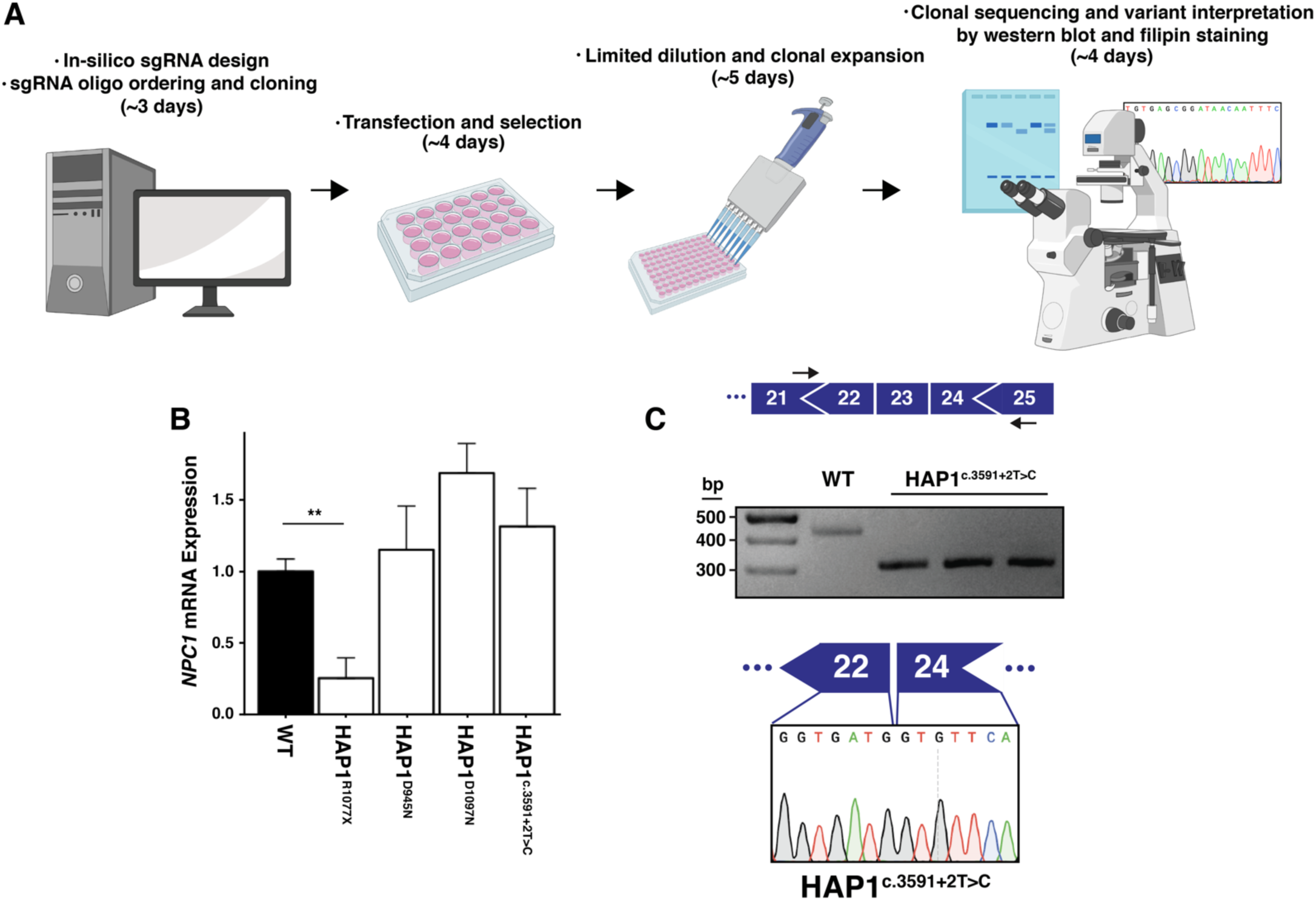
*NPC1* expression in NPC variant cell models. (A) Schematic overview of the process of cell model generation and characterization. (B) Expression of *NPC1* mRNA is significantly decreased in the NPC1 p.R1077X cell model (n = 3, p = 0.002 by two-tailed *t*-test) but unchanged in all other variants assayed. (C) Top, diagram illustrating PCR assay used to analyze splicing. Middle, PCR amplification results in a shorter amplicon in *NPC1* c.3591+2T>C Cells compared to wildtype. Bottom, sequencing chromatogram from *NPC1* c.3591+2T>C cells showing exclusion of exon 23.

### Haploid cell models of NPC1 variants allow for variant characterization and clinical interpretation

The majority of NPC patients are compound heterozygotes and often carry at least one private mutation (Park et al. 2003; Fernandez-Valero et al. 2005). As a consequence, it remains challenging to attribute a specific molecular mechanism of disease to an individual *NPC1* variant. Using our haploid models, we sought to characterize the nineteen aforementioned *NPC1* variants. First, we assayed the expression of *NPC1* mRNA in four of the cell models – NPC1 p.D945N, NPC1 p.R1077X, NPC1 p.D1097N and *NPC1* c.3591+2T>C – where variant interpretation has been previously documented. While there was a trend towards increased *NPC1* mRNA expression in NPC1 p.D945N, NPC1 p.D1097N and *NPC1* c.3591+2T>C compared to wildtype, no measurement reached significance (Fig. 2B). This aligns with previously reported data from NPC patient fibroblasts, where select missense mutations and in-frame deletion mutations have been shown to result in modestly increased *NPC1* mRNA expression (Gelsthorpe et al. 2008; Yamamoto et al. 2004). In the NPC1 p.R1077X cell model, however, there was a significant reduction in *NPC1* mRNA expression (25.3±14%, p = 0.002, n = 3, Fig. 2B). We suspect the reduction in *NPC1* mRNA expression is the result of nonsense mediated decay due to the introduced premature stop codon, as previously reported for other nonsense variants in NPC and a variety of other genetic disorders (Macias-Vidal et al. 2009; Frischmeyer and Dietz 1999). The modelled splice-site mutation, *NPC1* c.3591+2T>C, is predicted to disrupt canonical splicing of the *NPC1* transcript. To assess splicing, we designed a cDNA-based PCR assay that amplified a region between exons 21 and 25 (Fig. 2C). In our assay, amplification of the *NPC1* c.3591+2T>C splice-site mutation model resulted in a band approximately 100bp shorter than the wildtype amplicon (Fig. 2C). By Sanger sequencing, we confirmed that the shorter amplicon was indeed the result of exon 23 (114bp in length) exclusion (Fig. 2C).

Next, by analyzing NPC1 protein expression via western blot, we found that mutant NPC1 expression varied in apparent molecular weight and level of expression. Six of the nineteen variants – NPC1 p.R4H, NPC1 p.F402P, NPC1 p.Y634=, NPC1 p. I635T, NPC1 p.V1158=, and NPC1 p.M1159T – ran with an equivalent molecular weight to wildtype NPC1 protein (Fig. 3). Each of these variants had similar NPC1 expression compared to wildtype, with exception of NPC1 p.F402P, which had a moderate reduction in protein expression. Seven of the nineteen variants – NPC1 p.E406G, NPC1 p.Y634C, NPC1 p.G765K, NPC1 p. L1060P, NPC1 p.I1061T, NPC1 p.D1097N, and *NPC1* c.3591+2T>C – ran as a single band at a lower molecular weight than wildtype NPC1 protein (Fig. 3) and exhibited reduced expression compared to wildtype. Of note, the reduction in protein expression found in the *NPC1* c.3591+2T>C implies that despite the exon 23 exclusion leaving the open-reading-frame intact, the protein is likely being targeted for degradation. Four of the nineteen variants – NPC1 p.F402S, NPC1 p.L785S, NPC1 p.D898N and NPC1 p.D945N – exhibited a reduction in total protein and ran as two bands, one equivalent to wildtype and the other equivalent to that found in the lower molecular weight mutants (Fig. 3). The distinct molecular weights found in a subset of the mutant variants modelled is consistent with findings in patient-derived fibroblasts and have been attributed to endoplasmic-reticulum-associated protein degradation and heterogeneous glycosylation (Gelsthorpe et al. 2008; Zampieri et al. 2012; Millat et al. 2001; Yamamoto et al. 2000; Nakasone et al. 2014; Watari et al. 1999). The NPC1 p.R1077X model, consistent with the reduced mRNA expression, exhibited a complete absence of NPC1 protein (Fig. 3). In preliminary assays, the NPC1 p.Y1267C model also displayed a total absence of NPC1 protein (Fig. 3). The amino acid change for this variant, however, occurs within immunogen sequence of the primary C-terminal antibody. Upon further analysis with an N-terminal antibody, we found that NPC1 p.Y1267C protein ran similarly to wildtype (Supplemental Fig. S4). Having observed different levels of expression and migration patterns in our models, we assayed cholesterol homeostasis in each of the cell lines (Fig. 4 and Supplemental Fig. S5). With the exception of NPC1 p.L785S, all variants that ran either as a single lower molecular weight band or a doublet demonstrated defective cholesterol trafficking indicated by the distinct foci of cholesterol deposits revealed by filipin staining. Interestingly, despite a marked reduction in NPC1 protein, the NPC1 p.L785S model appeared largely indistinguishable from wildtype HAP1 cells by filipin staining, with only a minor subset (∼5%) of cells exhibiting lysosomal cholesterol accumulation (Fig. 4). It is likely that this variant represents what has been well-documented in a minority of NPC patients – a variant biochemical phenotype. In these biochemical variants, filipin staining of patient-derived fibroblasts is less definitive (Vanier et al. 1991). Six of the nineteen variants – the two variants harbouring silent mutations, NPC1 p.Y634= and NPC1 p.V1158=, and the missense mutants NPC1 p.R4H, NPC1 p.I635T, NPC1 p.M1159T and NPC1 p.Y1267C – appeared comparable to wildtype (Supplemental Fig. S5).

**Figure 3.**
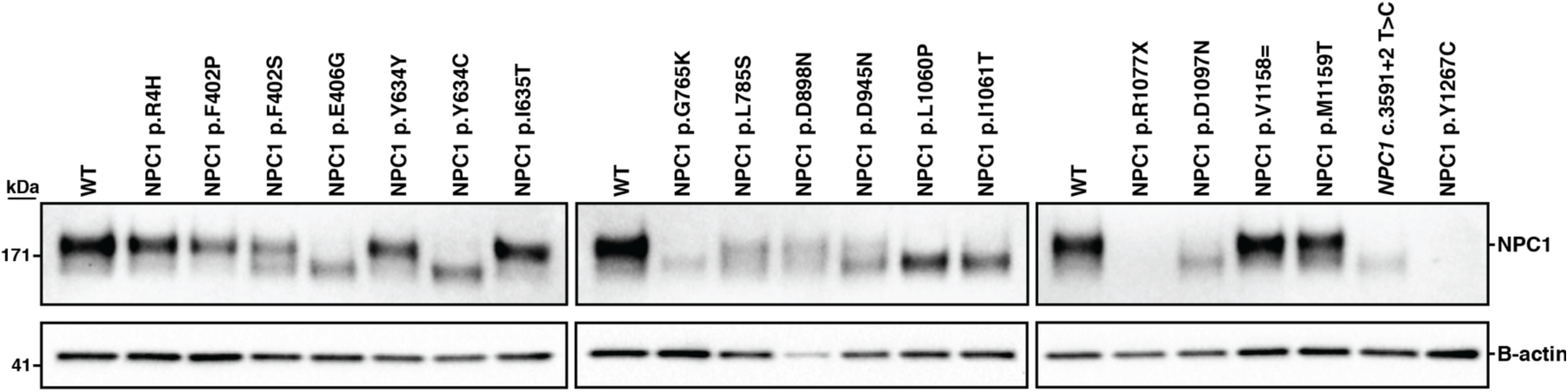
NPC1 expression varies across haploid cell models of Niemann-Pick disease type C. Expression of NPC1 protein was measured via western blot for all NPC1 variants modelled. B-actin was used a loading control.

**Figure 4.**
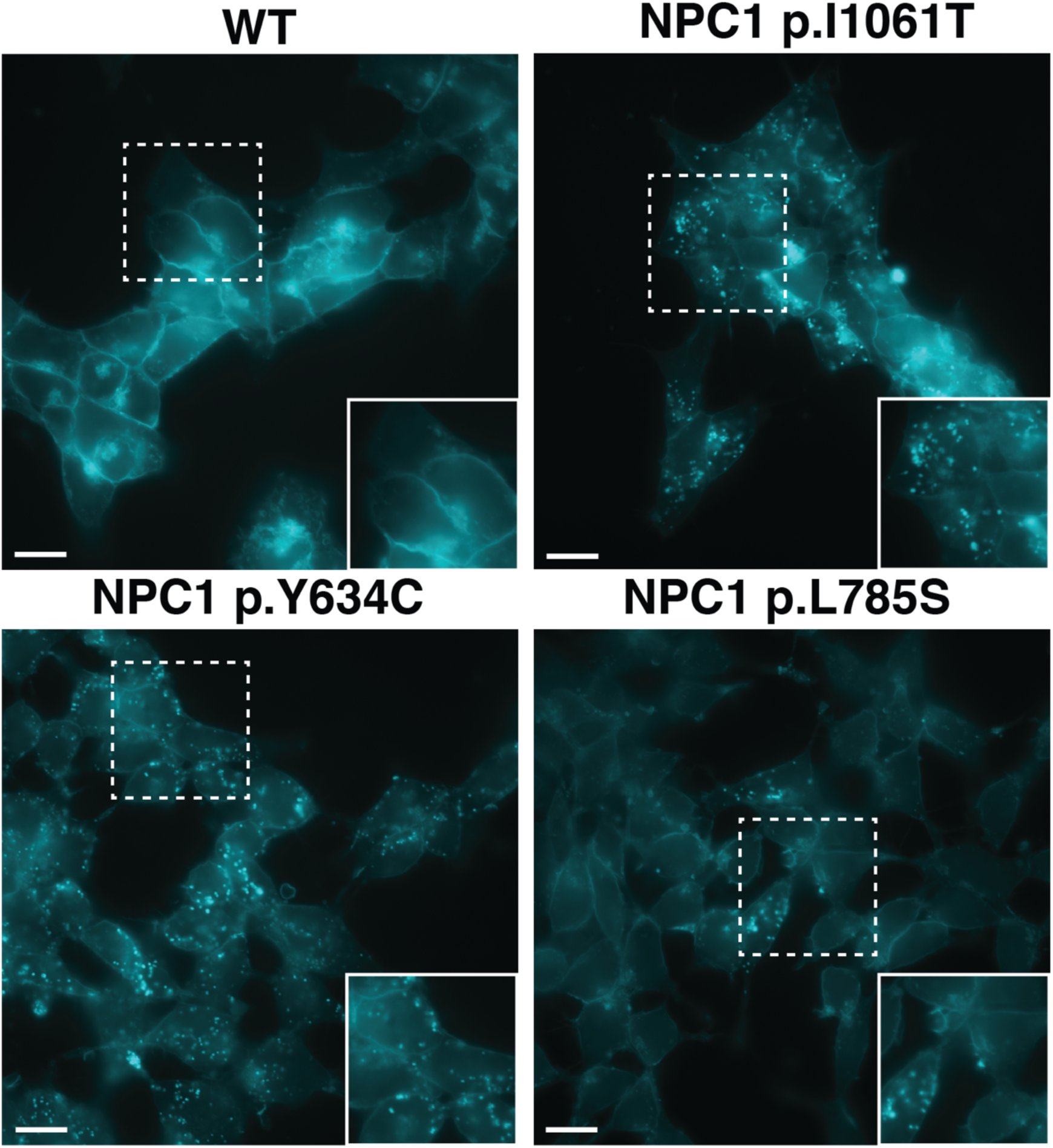
Filipin staining of haploid models of NPC1 variants. Wildtype HAP1 cells, top left, displayed no defect in cholesterol trafficking. Pathogenic variants, NPC1 p.I1061T and NPC1 p.Y634C, top right and bottom left respectively, exhibit distinct foci of filipin staining indicative of a defect in cholesterol trafficking. NPC1 p.L785S, bottom right, was found to exhibit a biochemical variant phenotype, with less definitive filipin staining. White dashed-bordered box has been enlarged twofold and inset at bottom right. Scale bars: 13μm.

Taken together, this data provides strong functional evidence for the clinical pathogenicity of each variant modelled, summarized in Table 1. Critically, our data resolve the clinical interpretation of three variants presently documented as variants of uncertain significance – NPC1 p.Y634C, NPC1 p.D898N and NPC1 p.D945N – indicating that each variant is pathogenic (Landrum et al. 2016). For the remaining variants, we have either confirmed existing clinical interpretations, or made the first interpretation of clinical significance. Together, these data demonstrate the utility of our haploid models of *NPC1* variants in the delineation of disease mechanisms and the interpretation of clinical variants.

## Discussion

Using CRISPR-Cas9-based genome editing, we have developed the first haploid cell model of Niemann-Pick disease type C. Our model recapitulates the primary biochemical and diagnostic phenotype found in patient-derived fibroblasts. While other human cells lines deficient in NPC1 expression have been reported (Zhao and Ridgway 2017; Tharkeshwar et al. 2017; Du, Lukmantara, and Yang 2017; Rodriguez-Pascau et al. 2012), these have all been diploid or aneuploid, without established uniform allelic disruption, and NPC1 deficiency confirmed by immunoblotting only. Using CRISPR-Cas9-mediated gene editing we obtained two unique NPC1-null cell models with indels resulting in verified coding frameshifts, and one NPC1-deficient cell model containing a six base pair in-frame deletion. Similar to NPC patient-derived fibroblasts, these models display a distinct defect in cholesterol trafficking, resulting in the accumulation of unesterified cholesterol.

To date, greater than 200 disease-causing mutations in *NPC1* have been reported. Accordingly, it is common for NPC patients to be compound heterozygotes, often harbouring at least one private mutation. The precise nature of a given mutation in *NPC1* can vary widely, including both missense and nonsense mutations, splice-site mutations, small duplication mutations, and indel mutations (Millat et al. 2001; Tarugi et al. 2002; Fernandez-Valero et al. 2005; Park et al. 2003). Despite the vast diversity found in the *NPC1* mutation spectrum, most research detailing the molecular mechanisms of NPC have been focused on one mutation, NPC1 p.I1061T (Gelsthorpe et al. 2008; Rauniyar et al. 2015; Schultz et al. 2018). As this allele accounts for between 15-20% of all NPC disease alleles (Davies and Ioannou 2000; Millat et al. 2001; Park et al. 2003), it is readily available in homozygous patient fibroblasts – a unique occurrence given the well-documented heterogeneity of *NPC1* mutations. A detailed understanding of the majority of *NPC1* variants using patient-derived fibroblasts, however, remains challenging as they are rarely found in isolation.

Here, we have developed a platform that allows for the rapid generation and analysis of *NPC1* variants. Specifically, our approach allows for the isolation and expansion of mutant cell clones in less than 14 days. Using CRISPR-Cas9-mediated base editing, we modelled nineteen unique *NPC1* variants, demonstrating the utility of the system in terms of variant characterization and clinical interpretation, and expanding our understanding of the genotype-phenotype relation in *NPC1*. The nineteen modelled variants demonstrate varied outcomes of *NPC1* mutations at the level of mRNA and protein, suggesting different mechanisms of pathogenicity. As such, our results emphasize the need for a mutation-by-mutation analysis of the *NPC1* gene, which could foreseeably lead to further insights into basic *NPC1* function and help identify unique therapeutic avenues.

Importantly, CRISPR-Cas9-mediated base editing in HAP1 cells is so efficient that, by screening only a moderate number of clones one can isolate multiple positive colonies. Recently, two studies have documented off-target editing events as a result of cytosine base editors (Zuo et al. 2019; Jin et al. 2019). Importantly, it was demonstrated that these off-target events were independent of sgRNA sequence and were enriched in highly transcribed regions. Accordingly, we carried out an analysis on three independent cell clones for each variant of interest, minimizing the potential that any observed phenotype to be off-target dependent. Furthermore, we performed RNA-sequencing on the parental HAP1 line and nine edited cell clones representing the three biological replicates from three of our modelled variants, NPC1 p.R1077X, NPC1 p.D1097N and *NPC1* c.3591+2T>C. Using these sequencing reads, we performed variant analysis evaluating the distribution of mutations throughout the cell clones. While we found between 9 to 20 mutations in each clone that were not present in the parental cell line, no single mutation was shared by multiple cell clones indicating that the observed cholesterol accumulation phenotype is independent of potential off-target base editing in the transcriptome (Supplemental Fig. S6A). Furthermore, of the mutations found there was no apparent bias towards C to T or G to A substitutions in clones generated using cytosine base editing and no apparent bias towards A to G or T to C substitutions in clones generated using adenine base editing (Supplemental Fig. S6B). This indicates the mutations observed are unlikely true off-targets, but likely a result of random mutation events.

The American College of Medical Genetics guidelines classify the results of functional assays as strong evidence for or against variant pathogenicity (Richards et al. 2015). We have shown that *NPC1* perturbation leads to a readily visualized defect in cholesterol trafficking in HAP1 cells. This makes haploid models of *NPC1* variants an ideal system for clinical variant interpretation. We demonstrated this by confirming the clinical interpretation of four documented *NPC1* variants, resolving the clinical interpretation of three *NPC1* variants of uncertain significance and establishing the clinical interpretation of twelve presently undocumented *NPC1* variants.

Given the overt phenotype induced by *NPC1* perturbation it can be envisioned that, if appropriately scaled, a CRISPR-Cas9-mediated base editing screen using HAP1 cells could serve as an effective platform for the clinical interpretation of a multitude of NPC variants. Presently, there are 1839 non-synonymous *NPC1* mutations documented in the GnomAD database (Lek et al. 2016), 56% of which, are the result of transitional nucleotide substitutions. In the present study, we demonstrated efficient targeting using multiple engineered SpCas9 variants (Table 1). Indeed, with the expanding list of available Cas9 enzymes, each with a unique PAM consensus sequence (Hu et al. 2018; Hua, Tao, and Zhu 2019; Yang et al. 2018), it is foreseeable that the vast majority of documented *NPC1* variants could be modelled using CRISPR-Cas9-mediated base editing. Furthermore, base editing systems are not strictly limited to transitional nucleotide substitutions, which further expands their utility in variant interpretation. While occurring in a less predictable manner, the targeted activation-induced deaminase (AID)-mediated mutagenesis system allows for all nucleotide substitutions of a cytosine (Ma et al. 2016) and the CRISPR-X system allows for all substitutions of both cytosines and guanines (Hess et al. 2016; Ma et al. 2016). More recently still, it was demonstrated that adenine base editors are capable of cytosine deamination, ultimately resulting in both cytosine to guanine or cytosine to thymine substitutions (Kim et al. 2019).

Presently, our approach is limited by the editing window of current base editing technologies, which is approximately five base pairs wide (Komor et al. 2016; Komor et al. 2017; Rees et al. 2017). While the percentage of total modellable mutations are limited by the currently permissible editing, efforts are underway to engineer enzymes with a much narrower editing window (Tan et al. 2019). Interestingly, we noted at least one editing event with nucleotide substitutions occurring well-outside of the predicted editing window for both SpCas9 cytosine and SpCas9 adenine base editors (Supplemental Fig. S3). While these events were in the minority, they suggest the boundaries of the SpCas9 base editing window are not absolute. Accordingly, it may be possible to design enzymes with a shifted editing window, further expanding the catalog of targetable mutations.

In this study, using CRISPR-Cas9-mediated gene editing, we have generated and characterized three models of NPC1-deficiency, which represent the first human haploid cell models of NPC. We demonstrated that these models effectively capture the principle diagnostic readout of NPC. The sheer number of *NPC1* variants, combined with most patients being compound heterozygotes, presents a challenge to understanding the disease mechanisms underlying individual patient variants. To overcome this challenge, we sought to model unique *NPC1* variants in HAP1 cells. We demonstrated that this is readily achievable using CRISPR-Cas9-mediated base editing. Finally, we showed that our haploid models of *NPC1* variants allow for efficient variant characterization and clinical interpretation. While we have focused on NPC, it is worth of note that the largest class of known pathogenic mutations in humans are point mutations (Landrum et al. 2016; Landrum et al. 2014). Given the ease and efficiency of our approach, one can envision applying this strategy to a wide variety of genetic disorders, thus providing a platform for the establishment of detailed genotype-phenotype relationships across these many disorders.

## Materials and Methods

### Cell culture and transfection

HAP1 cells were obtained from Horizon Genomics (Cambridge, United Kingdom). HAP1 cells were cultured in Iscove’s modified Dulbecco’s medium (Gibco) supplemented with 10% fetal bovine serum, 1% L-glutamine and 1% Pen-Strep (Gibco). Patient-derived fibroblasts were cultured in DMEM (Gibco) supplemented with 10% fetal bovine serum, 1% L-glutamine and 1% Pen-Strep (Gibco). We seeded 500,000 HAP1 cells for transfection in 2mL of media in six-well plates using Lipofectamine 3000 (Thermo Fisher). Cells were transfected with 1250 ng of a Cas9 base editor expression vector, and 1250 ng of a sgRNA expression vector containing a puromycin resistance gene. The sgRNA expression vector was generated as follows: a puromycin resistance gene was PCR amplified from the pSpCas9(BB)-2A-Puro (PX459) V2.0 plasmid, which was a gift from Feng Zhang (Addgene plasmid # 62988), appending an AgeI cut site to the 5’ end of the amplicon. This amplicon was inserted into the pX600-AAV-CMV::NLS-SaCas9-NLS-3xHA-bGHpA plasmid, which was a gift from Feng Zhang (Addgene plasmid # 61592), replacing the SaCas9 coding region and resulting in a CMV-driven puromycin resistance gene. Using InFusion cloning (CloneTech) this puromycin expression cassette was inserted into an EcoRI linearized BPK1520 plasmid, which was a gift from Keith Joung (Addgene plasmid # 65777). The pLenti-FNLS-P2A-Puro plasmid used for a portion of the cytosine base editing experiments was a gift from Lukas Dow (Addgene plasmid # 110841). The pSI-Target-AID-NG plasmid used for the remaining cytosine base editing experiments was a gift from Osamu Nureki (Addgene plasmid # 119861). The pCMV_ABEmax_P2A_GFP, VRQR-ABEmax, NG-ABEmax plasmids used for adenine base editing experiments were gifts from David Liu (Addgene plasmid # 112101, 119811, 124163). To enrich for transfected cells, 24 hours post-transfection cells were subjected to 0.8 µg/mL of puromycin for 72 hours. Transfected cells were expanded for genomic DNA isolation and limited dilution as previously described (Essletzbichler et al. 2014).

### Genomic DNA isolation and PCR

Genomic DNA was isolated using the DNeasy Blood and Tissue Kit (Qiagen) according to the manufacturer’s protocol. PCR was performed using DreamTaq Polymerase (Thermo Fisher Scientific) according to the manufacturer’s protocol.

### Estimation of genome editing

PCR amplification from the genomic DNA of a bulk population of edited cells centered on the predicted editing site was performed. These amplicons were PCR purified using QIAquick PCR Purification Kit following the manufacturer’s protocol (Qiagen) and Sanger sequenced using the forward primer. To test guide efficiency, the Sanger sequencing files from unedited and edited cells were used as an input into the online sequence trace decomposition software, TIDE (Brinkman et al. 2014). To ascertain base editing percentage, the Sanger sequencing ab1 files were input into the online base editing analysis software, editR (G. et al. 2018; Kluesner et al. 2017).

### Protein isolation and western blot analysis

HAP1 cells were trypsinized from their well, pelleted, and washed three times with 1X PBS (Gibco). Protein was isolated from HAP1 cells by resuspending in 150uL of a one-to-one solution of RIPA homogenizing buffer (50 mM Tris HCl pH 7.4, 150 nM NaCl, 1-mM EDTA) and RIPA double-detergent buffer (2% deoxycholate, 2% NP40, 2% Triton X-100 in RIPA homogenizing buffer) supplemented with a protease-inhibitor cocktail (Roche). Cells were subsequently incubated on ice for 30 minutes. Cells were then centrifuged at 12000G for 15 minutes at 4**°**C and the supernatant was collected and stored at −80**°**C. Whole protein concentration was measured using Pierce BCA Protein Assay Kit according to the manufacturer’s protocol (Thermo Fisher Scientific). SDS-Page separation was completed by running 2 μg of total protein on a NuPAGE™ 3-8% Tris-Acetate gel (Thermo Fisher Scientific). Next, proteins were transferred to a nitrocellulose membrane using the iBlot 2 transfer apparatus (Thermo Fisher Scientific). A 5% milk solution in 1X TBST was used for blocking for 1 hour at room temperature. The membrane was then incubated with the NPC1 primary antibody (Abcam: ab106534 in main body figures or Novus Biologicals H00004864-M02 in Supplemental Fig. S4) at 4°C overnight. Primary antibody solution was removed, and the membrane was washed three times with 1X TBST. This was followed by a 1-hour Incubation at room temperature with horseradish peroxidase conjugated goat anti-rabbit IgG (Abcam). After three washes with 1X TBST, signal detection was achieved using SuperSignal™ West Femto Maximum Sensitivity Substrate (Thermo Fisher Scientific) according to the manufacturer’s protocol.

### Filipin staining

Coverslips were incubated for one hour at 37°C in 1mL of of 1:30 solution of Collagen I Rat Protein (Thermo Fisher Scientific) to 1X PBS. The collagen solution was then aspirated and 250,000 HAP1 cells were seeded onto the coverslips in 12-well plate 24 hours prior to staining. Cells were washed three times with 1X PBS, then fixed in 4% paraformaldehyde in 1X PBS for 30 minutes at room temperature. After fixation, cells were washed three times in 1X PBS, then stained with 50μg/mL of filipin III (Sigma) for one hour at room temperature. Coverslips were then washed three times in 1X PBS and mounted onto a slide using ProLong™ Gold Antifade Mountant (Thermo Fisher Scientific). Cells were visualized using the Zeiss Axiovert 200M epifluorescent microscope and images were captured with a Hamamatsu C4742-80-12AG camera.

### RNA isolation and quantitative PCR

Cells were harvested, washed three times with 1X PBS, pelleted, then resuspended in TRIzol™ Reagent (Thermo Fisher Scientific). Isolation of mRNA was isolated following the manufacturer’s protocol. Next, 500μg of mRNA was reverse transcribed using SuperScript™ III Reverse Transcriptase (Thermo Fisher Scientific) following the manufacturer’s protocol. Quantitative PCR utilizing Fast SYBR green Master Mix (Qiagen) on a Step One Plus Real Time PCR (Applied Biosystems) was performed. *NPC1* expression was analyzed by amplification using a forward primer spanning the junction of exon 12 and 13, and a reverse primer specific to exon 13. Primers against endogenous GAPDH were used as an internal control. ΔΔCt was analyzed to assess fold changes between edited and unedited samples.

### RNA-Sequencing and off-target analysis

RNA sequencing was performed by the Centre for Applied Genetics in Toronto using the Illumina HiSeq 2500 system, producing 120-bp paired-end reads. Raw transcript reads were aligned to the GRCh38 human genome using HISAT2. The Picard program (v2.21.1) was used to mark duplicative and sort reads. The Genome Analysis Toolkit (GATK v4.1.3.0) was used to the to split reads that contained Ns in their CIGAR string (McKenna et al. 2010). Variant calling was conducted with both FreeBayes (v1.3.1) and LoFreq (v2.1.3) independently (Wilm et al. 2012; Garrison and Marth 2012). Only variants called by both software programs were considered true variants. Any variants with a read depth of less than four, or an allele frequency less than 0.4, were filtered from the final list of unique mutant variants. The exon coordinates of GRCh38 version 86 were retrieved from the Ensembl database (Hunt et al. 2018) and any called variants falling outside of these coordinates were excluded. The final list of variants was manually inspected using IGV software (Thorvaldsdottir, Robinson, and Mesirov 2013).

### Statistical analyses

All graphs were plotted as the mean with error bars indicating standard error. Differences between groups was assessed by two-tailed Student’s *t*-test. P-values less than 0.05 were considered statistically significant.

### Data Access

All sgRNA and primer sequences are available in Supplemental Table S2. RNA-sequencing reads have been submitted to the Sequence Read Archive.

## Acknowledgments

We would like to thank Niemann-Pick Canada and the Marcogliese Family Foundation for their support and commitment to our research into Niemann-Pick disease type C. We are grateful to Ebony Thompson, Sonia Evagelou and Kyle Lindsay for their technical assistance. Figure 2A was created with Biorender.com.

## Competing Interests

The authors declare no competing or financial interests.

## Author Contributions

Conceptualization: S.E., E.A.I., R.D.C. Methodology: S.E., R.A.B., T.M.I.B., E.M., L.Z., E.A.I., R.D.C. Formal Analysis: S.E., R.A.B., T.M.I.B., E.A.I., R.D.C. Investigation: S.E., R.A.B., T.M.I.B., E.M., E.A.I., R.D.C. Resources: E.A.I., R.D.C. Data curation: S.E., R.A.B., T.M.I.B., E.A.I., R.D.C. Writing – original draft preparation: S.E. Writing – review and editing: S.E., R.A.B., T.M.I.B., E.M., L.Z., E.A.I., R.D.C. Supervision: E.A.I., R.D.C. Project Administration: E.A.I., R.D.C. Funding Acquisition: S.E., E.A.I., R.D.C.

## Funding

This work was supported by the Rare Disease Foundation and the BC Children’s Hospital Foundation, [2304 to S.E], The Hospital for Sick Children, [Restracomp scholarship to S.E], Niemann Pick Canada and the Marcogliese Family Foundation.

## References

Brinkman, E. K., T. Chen, M. Amendola, and B. van Steensel. 2014. ‘Easy quantitative assessment of genome editing by sequence trace decomposition’, Nucleic Acids Res, 42: e168.

Carette, J. E., M. Raaben, A. C. Wong, A. S. Herbert, G. Obernosterer, N. Mulherkar, A. Kuehne, P. J. Kranzusch, A. M. Griffin, G. Ruthel, P. Dal Cin, J. M. Dye, S. P. Whelan, K. Chandran, and T. R. Brummelkamp. 2011. ‘Ebola virus entry requires the cholesterol transporter Niemann-Pick C1’, Nature, 477: 340–3.

Cho, S. W., S. Kim, J. M. Kim, and J. S. Kim. 2013. ‘Targeted genome engineering in human cells with the Cas9 RNA-guided endonuclease’, Nat Biotechnol, 31: 230–2.

Davies, J. P., and Y. A. Ioannou. 2000. ‘Topological analysis of Niemann-Pick C1 protein reveals that the membrane orientation of the putative sterol-sensing domain is identical to those of 3-hydroxy-3-methylglutaryl-CoA reductase and sterol regulatory element binding protein cleavage-activating protein’, J Biol Chem, 275: 24367–74.

Du, X., I. Lukmantara, and H. Yang. 2017. ‘CRISPR/Cas9-Mediated Generation of Niemann-Pick C1 Knockout Cell Line’, Methods Mol Biol, 1583: 73–83.

Essletzbichler, P., T. Konopka, F. Santoro, D. Chen, B. V. Gapp, R. Kralovics, T. R. Brummelkamp, S. M. Nijman, and T. Burckstummer. 2014. ‘Megabase-scale deletion using CRISPR/Cas9 to generate a fully haploid human cell line’, Genome Res, 24: 2059–65.

Fernandez-Valero, E. M., A. Ballart, C. Iturriaga, M. Lluch, J. Macias, M. T. Vanier, M. Pineda, and M. J. Coll. 2005. ‘Identification of 25 new mutations in 40 unrelated Spanish Niemann-Pick type C patients: genotype-phenotype correlations’, Clin Genet, 68: 245–54.

Findlay, G. M., R. M. Daza, B. Martin, M. D. Zhang, A. P. Leith, M. Gasperini, J. D. Janizek, X. Huang, L. M. Starita, and J. Shendure. 2018. ‘Accurate classification of BRCA1 variants with saturation genome editing’, Nature, 562: 217-+.

Frischmeyer, P. A., and H. C. Dietz. 1999. ‘Nonsense-mediated mRNA decay in health and disease’, Hum Mol Genet, 8: 1893–900.

G. Kluesner Mitchell, Nedveck Derek A., Lahr Walker S., Garbe John R., Abrahante Juan E., Webber Beau R., and Moriarity Branden S. 2018. ‘EditR: A Method to Quantify Base Editing from Sanger Sequencing’, The CRISPR Journal, 1: 239–50.

Garrison, Erik, and Gabor Marth. 2012. ‘Haplotype-based variant detection from short-read sequencing’, 1207.3907.

Gaudelli, N. M., A. C. Komor, H. A. Rees, M. S. Packer, A. H. Badran, D. I. Bryson, and D. R. Liu. 2017. ‘Programmable base editing of A.T to G.C in genomic DNA without DNA cleavage’, Nature, 551: 464-+.

Gelsthorpe, M. E., N. Baumann, E. Millard, S. E. Gale, S. J. Langmade, J. E. Schaffer, and D. S. Ory. 2008. ‘Niemann-Pick type C1 I1061T mutant encodes a functional protein that is selected for endoplasmic reticulum-associated degradation due to protein misfolding’, J Biol Chem, 283: 8229–36.

Greer, W. L., M. J. Dobson, G. S. Girouard, D. M. Byers, D. C. Riddell, and P. E. Neumann. 1999. ‘Mutations in NPC1 highlight a conserved NPC1-specific cysteine-rich domain’, American Journal of Human Genetics, 65: 1252–60.

Hess, G. T., L. Fresard, K. Han, C. H. Lee, A. Li, K. A. Cimprich, S. B. Montgomery, and M. C. Bassik. 2016. ‘Directed evolution using dCas9-targeted somatic hypermutation in mammalian cells’, Nat Methods, 13: 1036–42.

Hu, J. H., S. M. Miller, M. H. Geurts, W. Tang, L. Chen, N. Sun, C. M. Zeina, X. Gao, H. A. Rees, Z. Lin, and D. R. Liu. 2018. ‘Evolved Cas9 variants with broad PAM compatibility and high DNA specificity’, Nature, 556: 57–63.

Hua, K., X. Tao, and J. K. Zhu. 2019. ‘Expanding the base editing scope in rice by using Cas9 variants’, Plant Biotechnol J, 17: 499–504.

Huang, T. P., K. T. Zhao, S. M. Miller, N. M. Gaudelli, B. L. Oakes, C. Fellmann, D. F. Savage, and D. R. Liu. 2019. ‘Circularly permuted and PAM-modified Cas9 variants broaden the targeting scope of base editors’, Nat Biotechnol, 37: 626–31.

Hunt, S. E., W. McLaren, L. Gil, A. Thormann, H. Schuilenburg, D. Sheppard, A. Parton, I. M. Armean, S. J. Trevanion, P. Flicek, and F. Cunningham. 2018. ‘Ensembl variation resources’, Database (Oxford), 2018.

Jin, S., Y. Zong, Q. Gao, Z. Zhu, Y. Wang, P. Qin, C. Liang, D. Wang, J. L. Qiu, F. Zhang, and C. Gao. 2019. ‘Cytosine, but not adenine, base editors induce genome-wide off-target mutations in rice’, Science, 364: 292–95.

Jinek, M., K. Chylinski, I. Fonfara, M. Hauer, J. A. Doudna, and E. Charpentier. 2012. ‘A programmable dual-RNA-guided DNA endonuclease in adaptive bacterial immunity’, Science, 337: 816–21.

Kim, H. S., Y. K. Jeong, J. K. Hur, J. S. Kim, and S. Bae. 2019. ‘Adenine base editors catalyze cytosine conversions in human cells’, Nat Biotechnol, 37: 1145–48.

Kluesner, Mitchell, Derek Nedveck, Walker Lahr, John Garbe, Juan Abrahante, Beau Webber, and Branden Moriarity. 2017. ‘EditR: A novel base editing quantification software using Sanger sequencing’, bioRxiv: 213496.

Koblan, L. W., J. L. Doman, C. Wilson, J. M. Levy, T. Tay, G. A. Newby, J. P. Maianti, A. Raguram, and D. R. Liu. 2018. ‘Improving cytidine and adenine base editors by expression optimization and ancestral reconstruction’, Nat Biotechnol, 36: 843–46.

Komor, A. C., Y. B. Kim, M. S. Packer, J. A. Zuris, and D. R. Liu. 2016. ‘Programmable editing of a target base in genomic DNA without double-stranded DNA cleavage’, Nature, 533: 420-+.

Komor, A. C., K. T. Zhao, M. S. Packer, N. M. Gaudelli, A. L. Waterbury, L. W. Koblan, Y. B. Kim, A. H. Badran, and D. R. Liu. 2017. ‘Improved base excision repair inhibition and bacteriophage Mu Gam protein yields C:G-to-T:A base editors with higher efficiency and product purity’, Sci Adv, 3: eaao4774.

Landrum, M. J., J. M. Lee, M. Benson, G. Brown, C. Chao, S. Chitipiralla, B. Gu, J. Hart, D. Hoffman, J. Hoover, W. Jang, K. Katz, M. Ovetsky, G. Riley, A. Sethi, R. Tully, R. Villamarin-Salomon, W. Rubinstein, and D. R. Maglott. 2016. ‘ClinVar: public archive of interpretations of clinically relevant variants’, Nucleic Acids Res, 44: D862–8.

Landrum, M. J., J. M. Lee, G. R. Riley, W. Jang, W. S. Rubinstein, D. M. Church, and D. R. Maglott. 2014. ‘ClinVar: public archive of relationships among sequence variation and human phenotype’, Nucleic Acids Research, 42: D980–D85.

Lek, M., K. J. Karczewski, E. V. Minikel, K. E. Samocha, E. Banks, T. Fennell, A. H. O’Donnell-Luria, J. S. Ware, A. J. Hill, B. B. Cummings, T. Tukiainen, D. P. Birnbaum, J. A. Kosmicki, L. E. Duncan, K. Estrada, F. Zhao, J. Zou, E. Pierce-Hoffman, J. Berghout, D. N. Cooper, N. Deflaux, M. DePristo, R. Do, J. Flannick, M. Fromer, L. Gauthier, J. Goldstein, N. Gupta, D. Howrigan, A. Kiezun, M. I. Kurki, A. L. Moonshine, P. Natarajan, L. Orozco, G. M. Peloso, R. Poplin, M. A. Rivas, V. Ruano-Rubio, S. A. Rose, D. M. Ruderfer, K. Shakir, P. D. Stenson, C. Stevens, B. P. Thomas, G. Tiao, M. T. Tusie-Luna, B. Weisburd, H. H. Won, D. Yu, D. M. Altshuler, D. Ardissino, M. Boehnke, J. Danesh, S. Donnelly, R. Elosua, J. C. Florez, S. B. Gabriel, G. Getz, S. J. Glatt, C. M. Hultman, S. Kathiresan, M. Laakso, S. McCarroll, M. I. McCarthy, D. McGovern, R. McPherson, B. M. Neale, A. Palotie, S. M. Purcell, D. Saleheen, J. M. Scharf, P. Sklar, P. F. Sullivan, J. Tuomilehto, M. T. Tsuang, H. C. Watkins, J. G. Wilson, M. J. Daly, D. G. MacArthur, and Consortium Exome Aggregation. 2016. ‘Analysis of protein-coding genetic variation in 60,706 humans’, Nature, 536: 285–91.

Li, X., F. Lu, M. N. Trinh, P. Schmiege, J. Seemann, J. Wang, and G. Blobel. 2017. ‘3.3 A structure of Niemann-Pick C1 protein reveals insights into the function of the C-terminal luminal domain in cholesterol transport’, Proc Natl Acad Sci U S A, 114: 9116–21.

Ma, Y., J. Zhang, W. Yin, Z. Zhang, Y. Song, and X. Chang. 2016. ‘Targeted AID-mediated mutagenesis (TAM) enables efficient genomic diversification in mammalian cells’, Nat Methods, 13: 1029–35.

Macias-Vidal, J., L. Gort, M. Lluch, M. Pineda, and M. J. Coll. 2009. ‘Nonsense-mediated mRNA decay process in nine alleles of Niemann-Pick type C patients from Spain’, Mol Genet Metab, 97: 60–4.

Mali, P., K. M. Esvelt, and G. M. Church. 2013. ‘Cas9 as a versatile tool for engineering biology’, Nat Methods, 10: 957–63.

Mali, P., L. Yang, K. M. Esvelt, J. Aach, M. Guell, J. E. DiCarlo, J. E. Norville, and G. M. Church. 2013. ‘RNA-guided human genome engineering via Cas9’, Science, 339: 823–6.

McKay Bounford, K., and P. Gissen. 2014. ‘Genetic and laboratory diagnostic approach in Niemann Pick disease type C’, J Neurol, 261 Suppl 2: S569–75.

McKenna, A., M. Hanna, E. Banks, A. Sivachenko, K. Cibulskis, A. Kernytsky, K. Garimella, D. Altshuler, S. Gabriel, M. Daly, and M. A. DePristo. 2010. ‘The Genome Analysis Toolkit: a MapReduce framework for analyzing next-generation DNA sequencing data’, Genome Res, 20: 1297–303.

Millat, G., C. Marcais, C. Tomasetto, K. Chikh, A. H. Fensom, K. Harzer, D. A. Wenger, K. Ohno, and M. T. Vanier. 2001. ‘Niemann-Pick C1 disease: Correlations between NPC1 mutations, levels of NPC1 protein, and phenotypes emphasize the functional significance of the putative sterol-sensing domain and of the cysteine-rich luminal loop’, American Journal of Human Genetics, 68: 1373–85.

Nakasone, N., Y. S. Nakamura, K. Higaki, N. Oumi, K. Ohno, and H. Ninomiya. 2014. ‘Endoplasmic reticulum-associated degradation of Niemann-Pick C1: evidence for the role of heat shock proteins and identification of lysine residues that accept ubiquitin’, J Biol Chem, 289: 19714–25.

Nishimasu, H., F. A. Ran, P. D. Hsu, S. Konermann, S. I. Shehata, N. Dohmae, R. Ishitani, F. Zhang, and O. Nureki. 2014. ‘Crystal structure of Cas9 in complex with guide RNA and target DNA’, Cell, 156: 935–49.

Nishimasu, H., X. Shi, S. Ishiguro, L. Gao, S. Hirano, S. Okazaki, T. Noda, O. O. Abudayyeh, J. S. Gootenberg, H. Mori, S. Oura, B. Holmes, M. Tanaka, M. Seki, H. Hirano, H. Aburatani, R. Ishitani, M. Ikawa, N. Yachie, F. Zhang, and O. Nureki. 2018. ‘Engineered CRISPR-Cas9 nuclease with expanded targeting space’, Science, 361: 1259–62.

Ory, D. S. 2000. ‘Niemann-Pick type C: a disorder of cellular cholesterol trafficking’, Biochim Biophys Acta, 1529: 331–9.

Park, W. D., J. F. O’Brien, P. A. Lundquist, D. L. Kraft, C. W. Vockley, P. S. Karnes, M. C. Patterson, and K. Snow. 2003. ‘Identification of 58 novel mutations in Niemann-Pick disease type C: correlation with biochemical phenotype and importance of PTC1-like domains in NPC1’, Hum Mutat, 22: 313–25.

Patterson, M. C., E. Mengel, F. A. Wijburg, A. Muller, B. Schwierin, H. Drevon, M. T. Vanier, and M. Pineda. 2013. ‘Disease and patient characteristics in NP-C patients: findings from an international disease registry’, Orphanet J Rare Dis, 8: 12.

Rauniyar, N., K. Subramanian, M. Lavallee-Adam, S. Martinez-Bartolome, W. E. Balch, and J. R. Yates, 3rd. 2015. ‘Quantitative Proteomics of Human Fibroblasts with I1061T Mutation in Niemann-Pick C1 (NPC1) Protein Provides Insights into the Disease Pathogenesis’, Mol Cell Proteomics, 14: 1734–49.

Rees, H. A., A. C. Komor, W. H. Yeh, J. Caetano-Lopes, M. Warman, A. S. B. Edge, and D. R. Liu. 2017. ‘Improving the DNA specificity and applicability of base editing through protein engineering and protein delivery’, Nat Commun, 8: 15790.

Rees, H. A., and D. R. Liu. 2018. ‘Base editing: precision chemistry on the genome and transcriptome of living cells’, Nat Rev Genet, 19: 770–88.

Richards, S., N. Aziz, S. Bale, D. Bick, S. Das, J. Gastier-Foster, W. W. Grody, M. Hegde, E. Lyon, E. Spector, K. Voelkerding, H. L. Rehm, and Acmg Laboratory Quality Assurance Committee. 2015. ‘Standards and guidelines for the interpretation of sequence variants: a joint consensus recommendation of the American College of Medical Genetics and Genomics and the Association for Molecular Pathology’, Genetics in Medicine, 17: 405–24.

Rodriguez-Pascau, L., M. J. Coll, J. Casas, L. Vilageliu, and D. Grinberg. 2012. ‘Generation of a human neuronal stable cell model for niemann-pick C disease by RNA interference’, JIMD Rep, 4: 29–37.

Schultz, M. L., K. L. Krus, S. Kaushik, D. Dang, R. Chopra, L. Qi, V. G. Shakkottai, A. M. Cuervo, and A. P. Lieberman. 2018. ‘Coordinate regulation of mutant NPC1 degradation by selective ER autophagy and MARCH6-dependent ERAD’, Nat Commun, 9: 3671.

Scott, C., and Y. A. Ioannou. 2004. ‘The NPC1 protein: structure implies function’, Biochim Biophys Acta, 1685: 8–13.

Tan, J., F. Zhang, D. Karcher, and R. Bock. 2019. ‘Engineering of high-precision base editors for site-specific single nucleotide replacement’, Nat Commun, 10: 439.

Tarugi, P., G. Ballarini, B. Bembi, C. Battisti, S. Palmeri, F. Panzani, E. Di Leo, C. Martini, A. Federico, and S. Calandra. 2002. ‘Niemann-Pick type C disease: mutations of NPC1 gene and evidence of abnormal expression of some mutant alleles in fibroblasts’, J Lipid Res, 43: 1908–19.

Tharkeshwar, A. K., J. Trekker, W. Vermeire, J. Pauwels, R. Sannerud, D. A. Priestman, D. Te Vruchte, K. Vints, P. Baatsen, J. P. Decuypere, H. Lu, S. Martin, P. Vangheluwe, J. V. Swinnen, L. Lagae, F. Impens, F. M. Platt, K. Gevaert, and W. Annaert. 2017. ‘A novel approach to analyze lysosomal dysfunctions through subcellular proteomics and lipidomics: the case of NPC1 deficiency’, Sci Rep, 7: 41408.

Thorvaldsdottir, H., J. T. Robinson, and J. P. Mesirov. 2013. ‘Integrative Genomics Viewer (IGV): high-performance genomics data visualization and exploration’, Brief Bioinform, 14: 178–92.

Vanier, M. T. 2010. ‘Niemann-Pick disease type C’, Orphanet J Rare Dis, 5: 16.

Vanier, M. T., C. Rodriguez-Lafrasse, R. Rousson, N. Gazzah, M. C. Juge, P. G. Pentchev, A. Revol, and P. Louisot. 1991. ‘Type C Niemann-Pick disease: spectrum of phenotypic variation in disruption of intracellular LDL-derived cholesterol processing’, Biochim Biophys Acta, 1096: 328–37.

Wassif, C. A., J. L. Cross, J. Iben, L. Sanchez-Pulido, A. Cougnoux, F. M. Platt, D. S. Ory, C. P. Ponting, J. E. Bailey-Wilson, L. G. Biesecker, and F. D. Porter. 2016. ‘High incidence of unrecognized visceral/neurological late-onset Niemann-Pick disease, type C1, predicted by analysis of massively parallel sequencing data sets’, Genetics in Medicine, 18: 41–48.

Watari, H., E. J. Blanchette-Mackie, N. K. Dwyer, M. Watari, E. B. Neufeld, S. Patel, P. G. Pentchev, and J. F. Strauss, 3rd. 1999. ‘Mutations in the leucine zipper motif and sterol-sensing domain inactivate the Niemann-Pick C1 glycoprotein’, J Biol Chem, 274: 21861–6.

Wilm, A., P. P. Aw, D. Bertrand, G. H. Yeo, S. H. Ong, C. H. Wong, C. C. Khor, R. Petric, M. L. Hibberd, and N. Nagarajan. 2012. ‘LoFreq: a sequence-quality aware, ultra-sensitive variant caller for uncovering cell-population heterogeneity from high-throughput sequencing datasets’, Nucleic Acids Res, 40: 11189–201.

Wojtanik, K. M., and L. Liscum. 2003. ‘The transport of low density lipoprotein-derived cholesterol to the plasma membrane is defective in NPC1 cells’, J Biol Chem, 278: 14850–6.

Yamamoto, T., J. H. Feng, K. Higaki, M. Taniguchi, E. Nanba, H. Ninomiya, and K. Ohno. 2004. ‘Increased NPC1 mRNA in skin fibroblasts from Niemann-Pick disease type C patients’, Brain Dev, 26: 245–50.

Yamamoto, T., H. Ninomiya, M. Matsumoto, Y. Ohta, E. Nanba, Y. Tsutsumi, K. Yamakawa, G. Millat, M. T. Vanier, P. G. Pentchev, and K. Ohno. 2000. ‘Genotype-phenotype relationship of Niemann-Pick disease type C: a possible correlation between clinical onset and levels of NPC1 protein in isolated skin fibroblasts’, J Med Genet, 37: 707–12.

Yang, L., X. Zhang, L. Wang, S. Yin, B. Zhu, L. Xie, Q. Duan, H. Hu, R. Zheng, Y. Wei, L. Peng, H. Han, J. Zhang, W. Qiu, H. Geng, S. Siwko, X. Zhang, M. Liu, and D. Li. 2018. ‘Increasing targeting scope of adenosine base editors in mouse and rat embryos through fusion of TadA deaminase with Cas9 variants’, Protein Cell, 9: 814–19.

Yasui, M., E. Suenaga, N. Koyama, C. Masutani, F. Hanaoka, P. Gruz, S. Shibutani, T. Nohmi, M. Hayashi, and M. Honma. 2008. ‘Miscoding properties of 2’-deoxyinosine, a nitric oxide-derived DNA Adduct, during translesion synthesis catalyzed by human DNA polymerases’, J Mol Biol, 377: 1015–23.

Zafra, M. P., E. M. Schatoff, A. Katti, M. Foronda, M. Breinig, A. Y. Schweitzer, A. Simon, T. Han, S. Goswami, E. Montgomery, J. Thibado, E. R. Kastenhuber, F. J. Sanchez-Rivera, J. Shi, C. R. Vakoc, S. W. Lowe, D. F. Tschaharganeh, and L. E. Dow. 2018. ‘Optimized base editors enable efficient editing in cells, organoids and mice’, Nat Biotechnol, 36: 888–93.

Zampieri, S., B. Bembi, N. Rosso, M. Filocamo, and A. Dardis. 2012. ‘Treatment of Human Fibroblasts Carrying NPC1 Missense Mutations with MG132 Leads to an Improvement of Intracellular Cholesterol Trafficking’, JIMD Rep, 2: 59–69.

Zhao, K., and N. D. Ridgway. 2017. ‘Oxysterol-Binding Protein-Related Protein 1L Regulates Cholesterol Egress from the Endo-Lysosomal System’, Cell Rep, 19: 1807–18.

Zuo, E., Y. Sun, W. Wei, T. Yuan, W. Ying, H. Sun, L. Yuan, L. M. Steinmetz, Y. Li, and H. Yang. 2019. ‘Cytosine base editor generates substantial off-target single-nucleotide variants in mouse embryos’, Science, 364: 289–92.

