## Supplementary figures and images for "Modelling Niemann-Pick disease type C in a human haploid cell line allows for patient variant characterization and clinical interpretation"

### Supplemental Figure S1

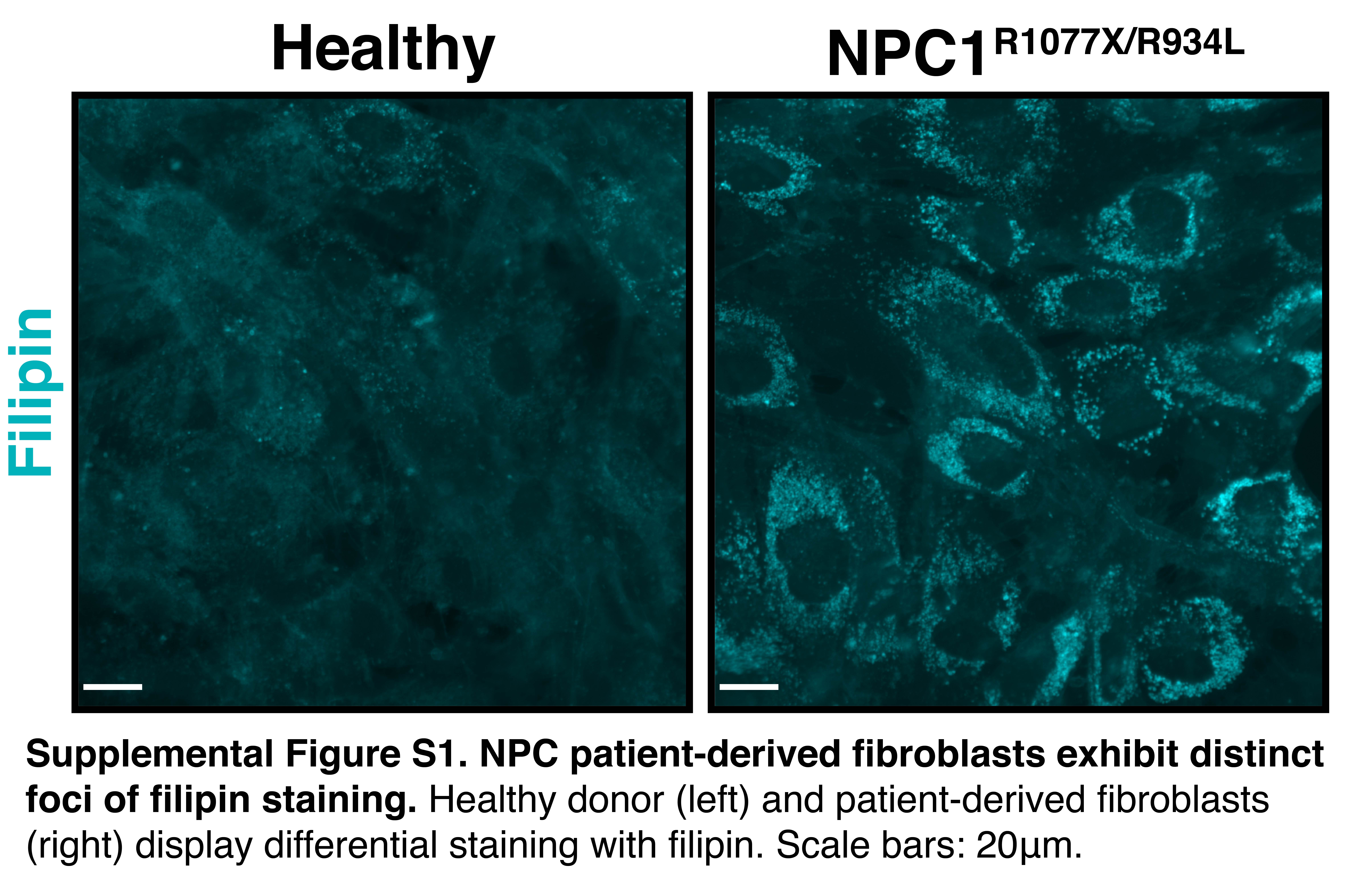

### Supplemental Figure S2

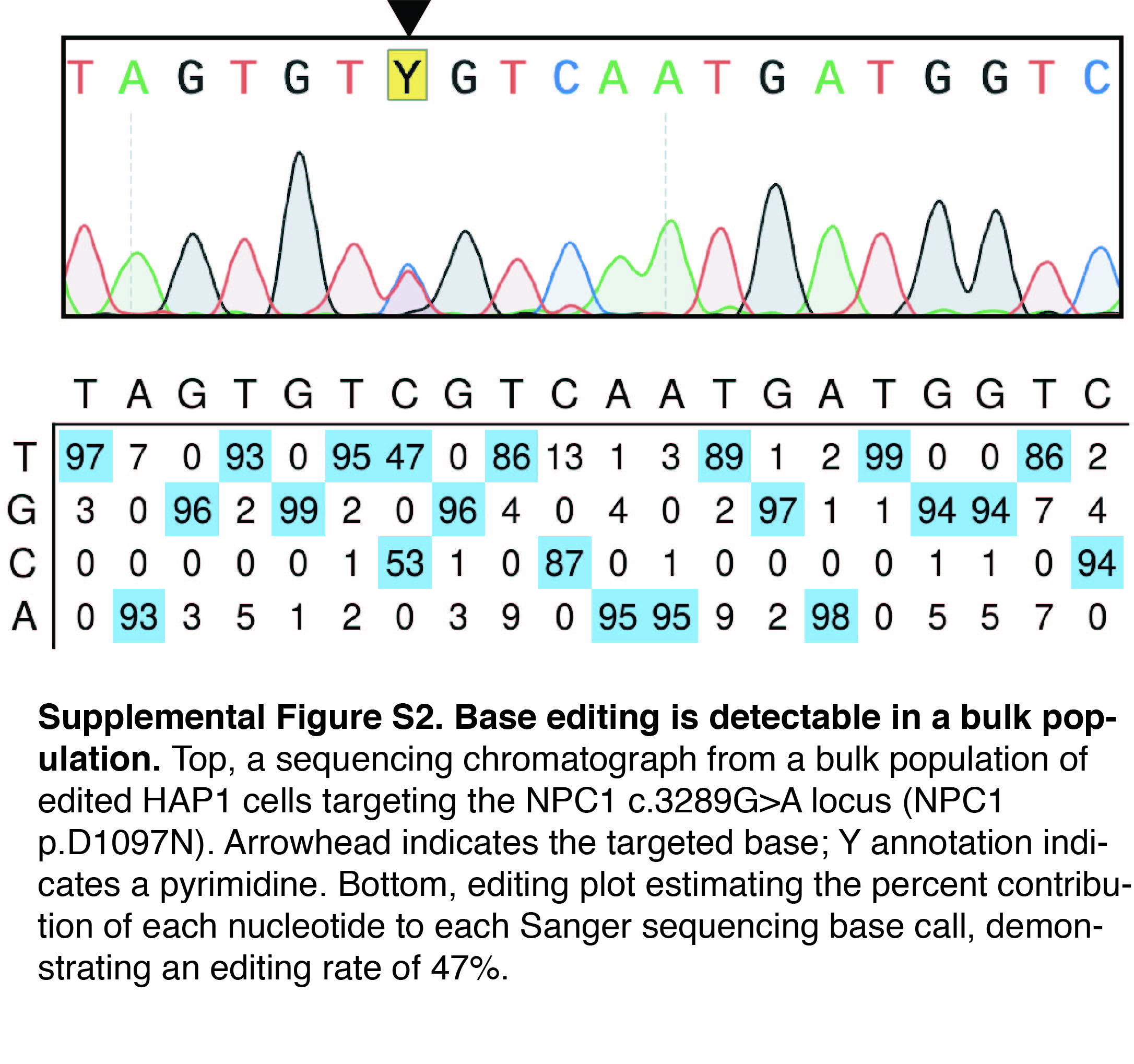

### Supplemental Figure S3

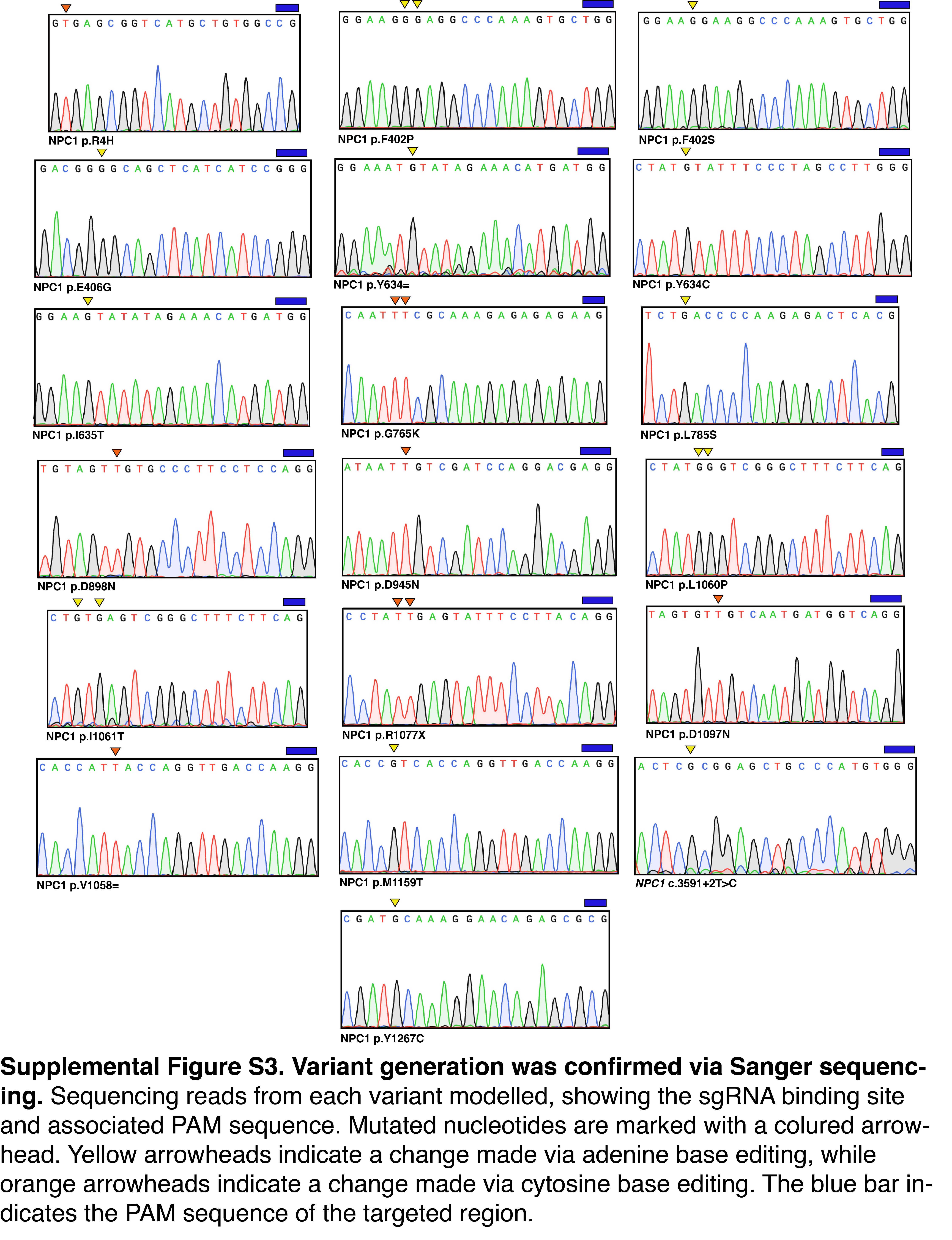

### Supplemental Figure S4

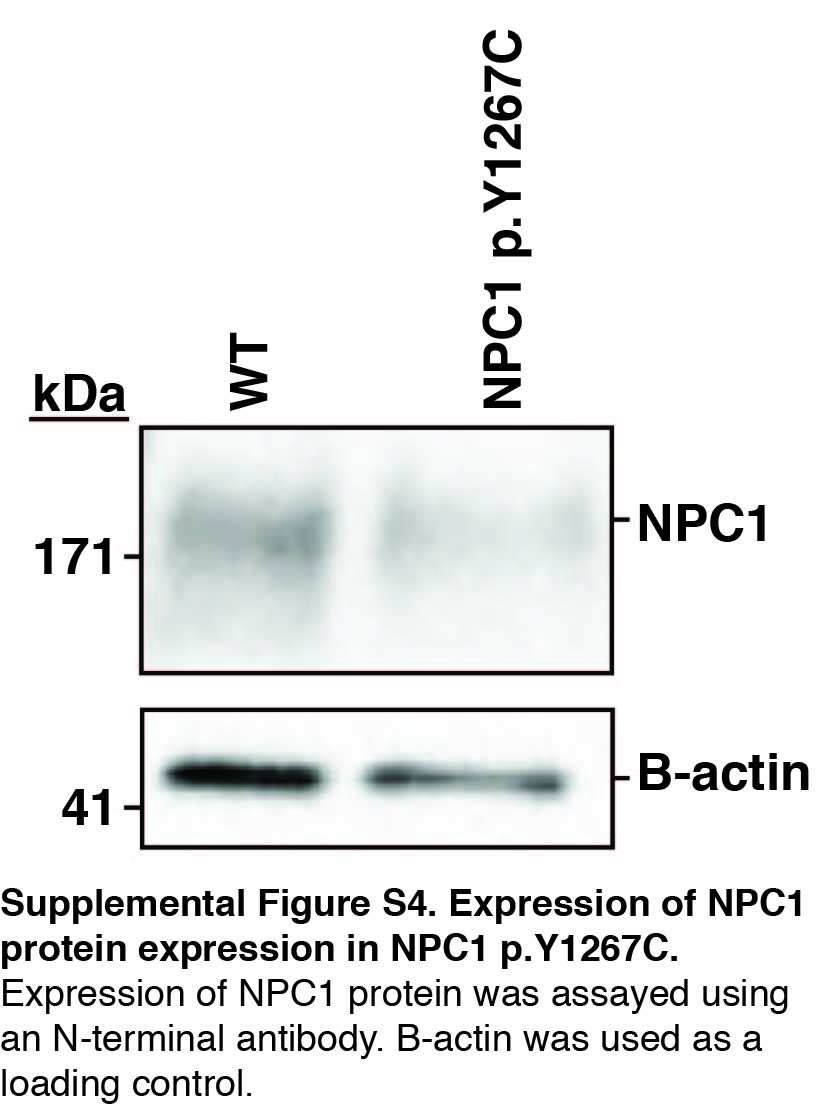

### Supplemental Figure S5

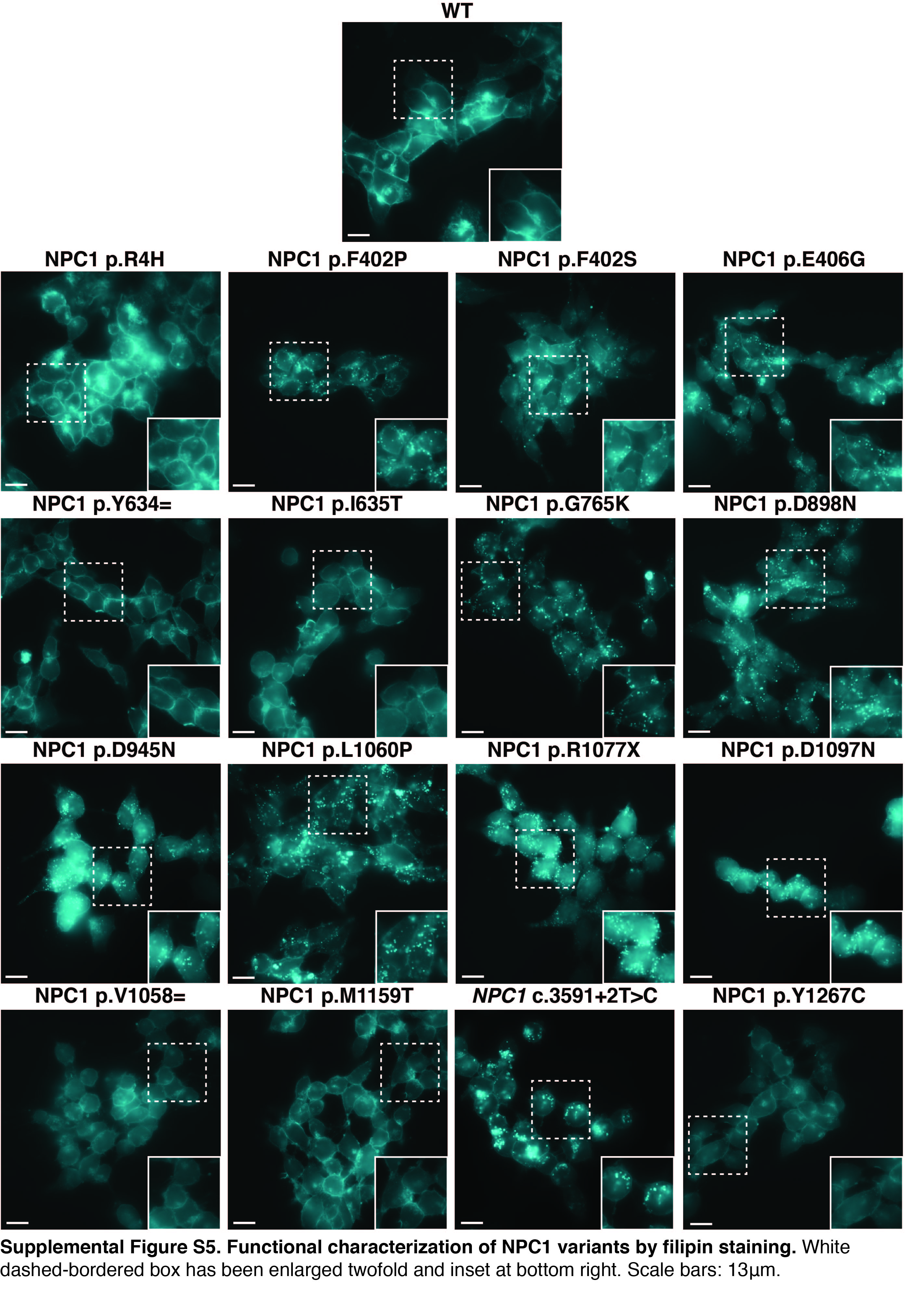
